# A novel clinical metaproteomics workflow enables bioinformatic analysis of host-microbe dynamics in disease

**DOI:** 10.1101/2023.11.21.568121

**Authors:** Katherine Do, Subina Mehta, Reid Wagner, Dechen Bhuming, Andrew T. Rajczewski, Amy P.N. Skubitz, James E. Johnson, Timothy J. Griffin, Pratik D. Jagtap

**Affiliations:** Department of Biochemistry, Molecular Biology and Biophysics, University of Minnesota, Minneapolis, MN, USA; Minnesota Supercomputing Institute, University of Minnesota, Minneapolis, MN, USA; Department of Laboratory Medicine and Pathology, University of Minnesota, Minneapolis, MN, USA

## Abstract

Clinical metaproteomics has the potential to offer insights into the host-microbiome interactions underlying diseases. However, the field faces challenges in characterizing microbial proteins found in clinical samples, which are usually present at low abundance relative to the host proteins. As a solution, we have developed an integrated workflow coupling mass spectrometry-based analysis with customized bioinformatic identification, quantification and prioritization of microbial and host proteins, enabling targeted assay development to investigate host-microbe dynamics in disease. The bioinformatics tools are implemented in the Galaxy ecosystem, offering the development and dissemination of complex bioinformatic workflows. The modular workflow integrates MetaNovo (to generate a reduced protein database), SearchGUI/PeptideShaker and MaxQuant (to generate peptide-spectral matches (PSMs) and quantification), PepQuery2 (to verify the quality of PSMs), and Unipept and MSstatsTMT (for taxonomy and functional annotation). We have utilized this workflow in diverse clinical samples, from the characterization of nasopharyngeal swab samples to bronchoalveolar lavage fluid. Here, we demonstrate its effectiveness via analysis of residual fluid from cervical swabs. The complete workflow, including training data and documentation, is available via the Galaxy Training Network, empowering non-expert researchers to utilize these powerful tools in their clinical studies.

## INTRODUCTION

Mass spectrometry (MS)-based metaproteomics enables the analysis of proteins expressed by microbial communities and can be applied to clinical samples to understand microorganism contributions to disease (1). Metaproteomics can provide insight into how the microbiome responds to a diseased condition by direct characterization of functional molecules (proteins) that are beyond the capabilities of metagenomics approaches, which mainly focus on taxonomic characterization (2). Moreover, clinical metaproteomics can provide insights into how the microbiome interacts with its host environment. However, one current challenge of metaproteomic analysis of clinical samples is that the high relative abundance of host (human) proteins can hamper the detection and identification of lower abundance microbial proteins. Moreover, identifying microbial peptides derived from tryptic digestion of isolated proteins involves searching tandem mass spectrometry (MS/MS) spectra against large sequence databases comprising all microbial proteomes present in the sample, decreasing sensitivity and increasing the potential for false positives (3). In addition, assigning taxonomy to the detected peptide sequences presents challenges due to the conservation of protein sequences across taxa (4). Assigning functions to detected proteins can also present challenges mainly due to a lack of confident annotation of encoded proteins (4, 5). Here, we offer a novel bioinformatics workflow that overcomes at least some of these challenges and enables effective metaproteomic analysis in clinical samples relevant to studies of disease.

To demonstrate the effectiveness of our clinical metaproteomics workflow, here we analyze MS/MS data from Pap test fluid (PTF) samples collected from ovarian cancer (OC) and non-OC patients. The bioinformatics workflow is accessible via the Galaxy ecosystem, which offers access to powerful bioinformatic tools for metaproteomic data analysis that facilitate the development and execution of complex workflows necessary for complete clinical metaproteomics (6). Galaxy is a free, browser-based, scalable platform that is maintained by a thriving community keeping it current to meet emerging needs in bioinformatics analysis across ‘omic domains (7, 8). New users can also access online and on-demand training resources, such as step-by-step instructions, access to workflows, and example datasets via Galaxy Training Network (GTN) resources (6, 9). We envision these collective bioinformatic resources will find use in many clinical studies, such as potential secondary infections during COVID-19 infection, and other broad research questions regarding host-microbe interactions underlying human disease underlying human disease (10). Such clinical metaproteomic studies offer the discovery of new microbe-host responses and interactions, as well as the potential to define peptide targets of interest for the development of targeted MS-based clinical assays.

## MATERIALS AND METHODS

### Sample processing to generate MS/MS spectra

For this workflow development, four example RAW files were used as input MS datasets. These RAW files were a subset of a total of forty PTF samples from twenty OC and twenty non-OC patients. Proteins from each sample were isolated, digested into peptides with trypsin, and labeled with a distinct Tandem Mass Tag (TMT)–11–plex tagging reagent. Each experimental group included one pooled reference sample labeled with a unique TMT tag that served as a common reference for comparison to each individual patient sample across all four separate experiments. The pooled samples were then separated by offline pH reversed–phase liquid chromatography (RPLC) and analyzed by liquid chromatography– tandem MS (LC–MS/MS), using a hybrid quadrupole–Orbitrap mass spectrometer to generate RAW MS/MS datasets. As required, the four example RAW files were converted into Mascot Generic Format (MGF) files for software compatibility, such as for SearchGUI/PeptideShaker searches in the Discovery Module. In the writing, the abbreviations “RAW” and “MGF” were included to denote the file format of the input MS datasets, and in the figures, the input MS datasets were represented by the same RAW dataset icons for simplification. As shown in **Figure 1**, the Galaxy bioinformatics workflow consists of five modules (Database Generation, Discovery, Verification, Quantification, and Data Interpretation). All modules are summarized in **Supplementary Table S1**, including details regarding inputs, outputs, and software tools. The complete workflow, datasets, and additional training resources are accessible via the Galaxy ecosystem and the GTN website (**Supplementary Tables S2, S3**).

**Figure 1.**
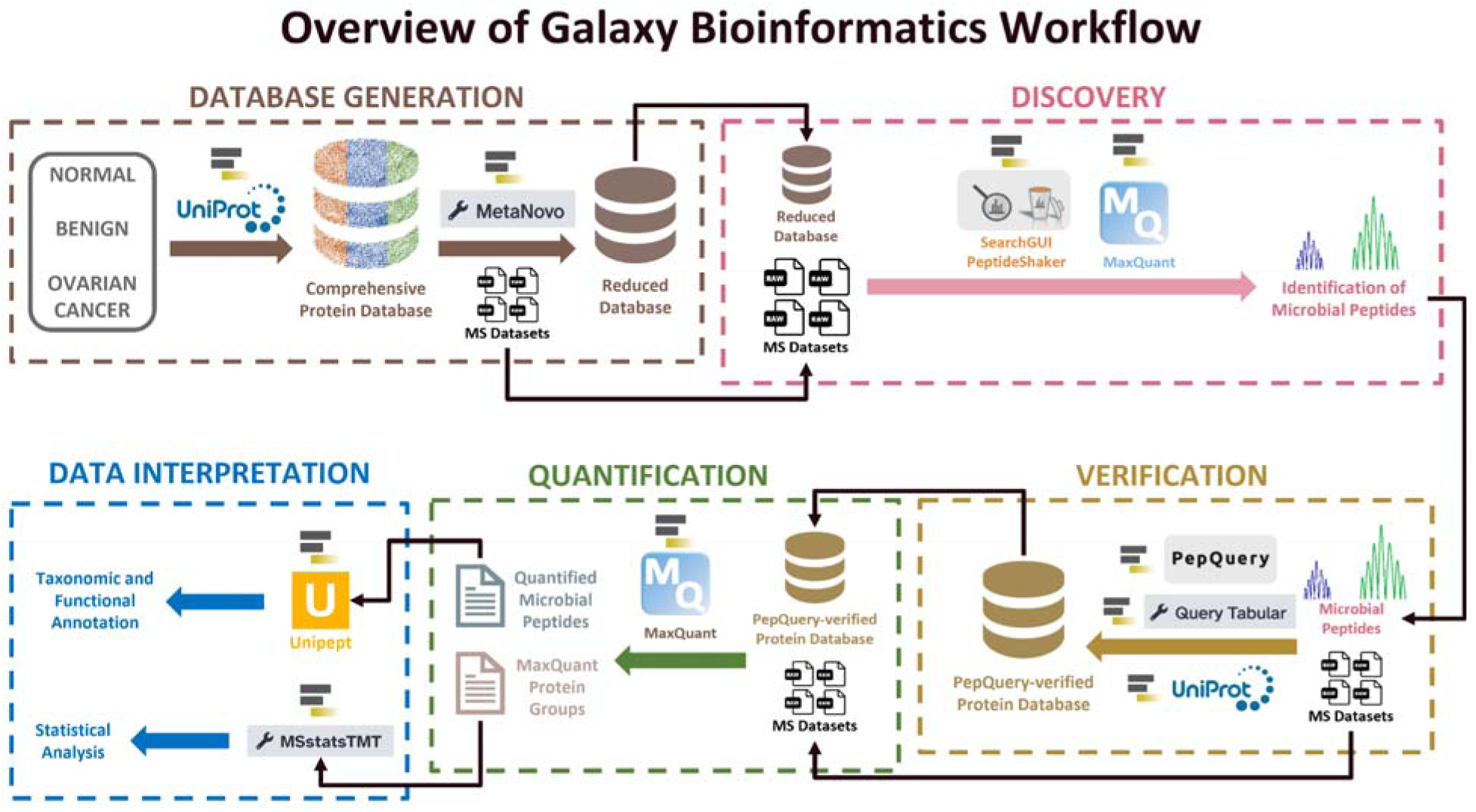
Overview of Galaxy Bioinformatic Workflow. This figure summarizes the workflow into five modules: Database Generation, Discovery, Verification, Quantification, and Data Interpretation.

### Database generation module for microbial peptide identification

To generate a comprehensive protein sequence database, a literature survey was conducted to obtain 118 taxonomic species of organisms that are commonly associated with the female reproductive tract (11).

Using this list of 118 taxonomic groups, a protein sequence FASTA database (3,383,217 sequences) was generated using the UniProt XML Downloader tool within the Galaxy framework. Additionally, Human SwissProt (reviewed-only; 20,408 sequences, as of September 2023) and contaminant (cRAP; 116 sequences) protein sequence databases were generated using the Protein Database Downloader tool.

The Species UniProt protein sequence database was then merged with the Human SwissProt (reviewed-only) and cRAP databases, using the FASTA Merge Files and Filter Unique Sequences tool to filter out duplicates and contaminants. This resulted in a comprehensive protein sequence database (2,595,745 sequences) (**Fig. 2**). This comprehensive database, along with the four MS datasets (MGF) generated by LC–MS/MS analysis of PTF samples, were inputs for the MetaNovo tool to generate a reduced databas (1,908 sequences) (12, 13). MetaNovo software uses outputs from DirecTag (*de novo* sequence tag matching tool) along with probabilistic optimization to generate customized protein sequence databases for target-decoy searches (13). The MetaNovo-generated database was then merged with the Human SwissProt (reviewed only) and cRAP databases to generate a compact database of 21,289 human and microbial sequences that were used for peptide identification.

**Figure 2.**
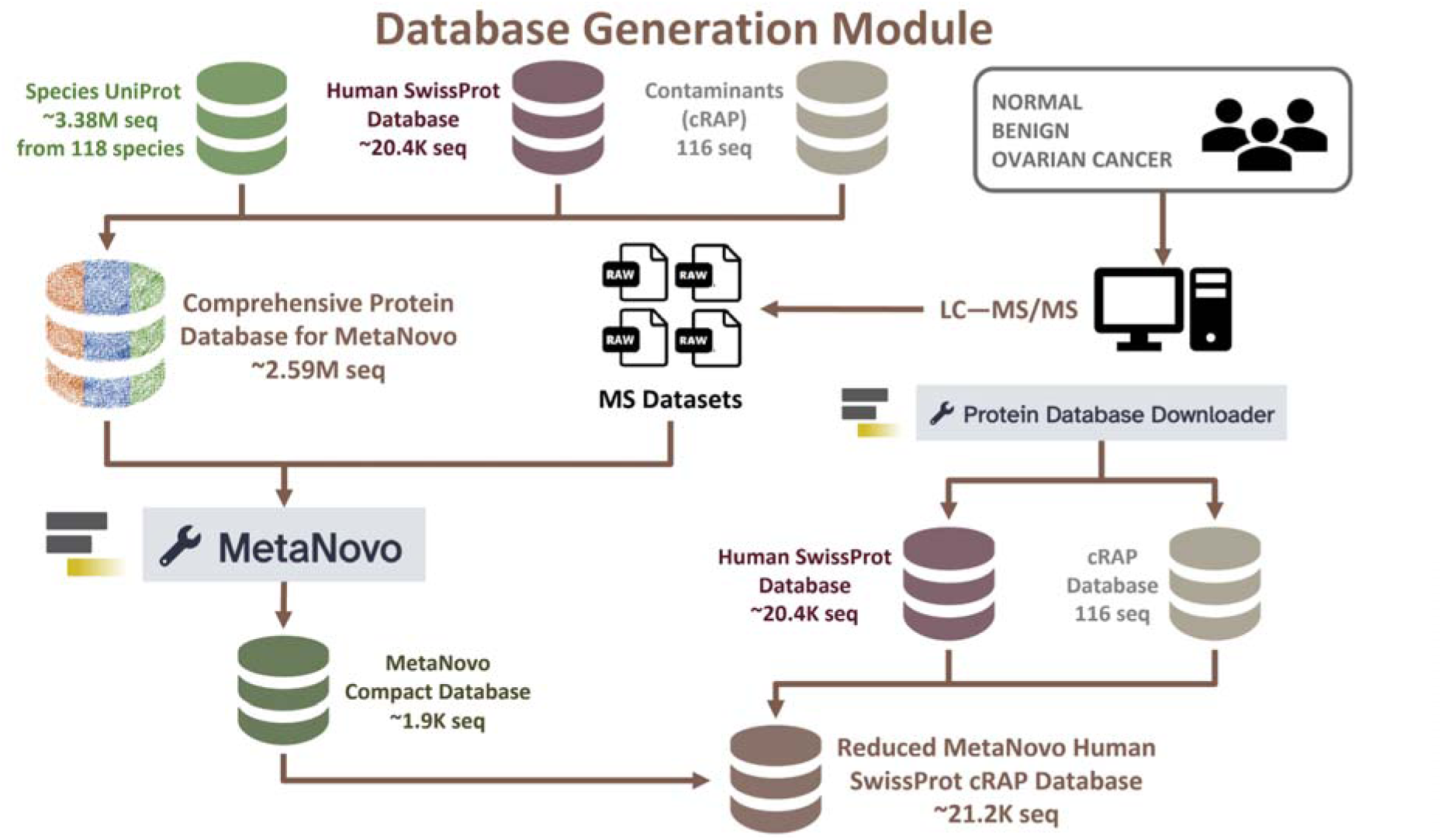
Database Generation Module. Overview of a large comprehensive database for input into MetaNovo and reduced database generation using MetaNovo.

### Discovery module using peptide identification programs

The four example MS datasets were searched against the compact database (21,289 sequences) to identify peptide sequences. Two peptide identification programs, SearchGUI/PeptideShaker and MaxQuant, were used for the searches (**Fig. 3; Supplementary Tables S4, S5**). For software compatibility, SearchGUI/PeptideShaker required the RAW files to be converted to MGF, using the msconvert tool, whereas MaxQuant is able to process the RAW files as is.

**Figure 3.**
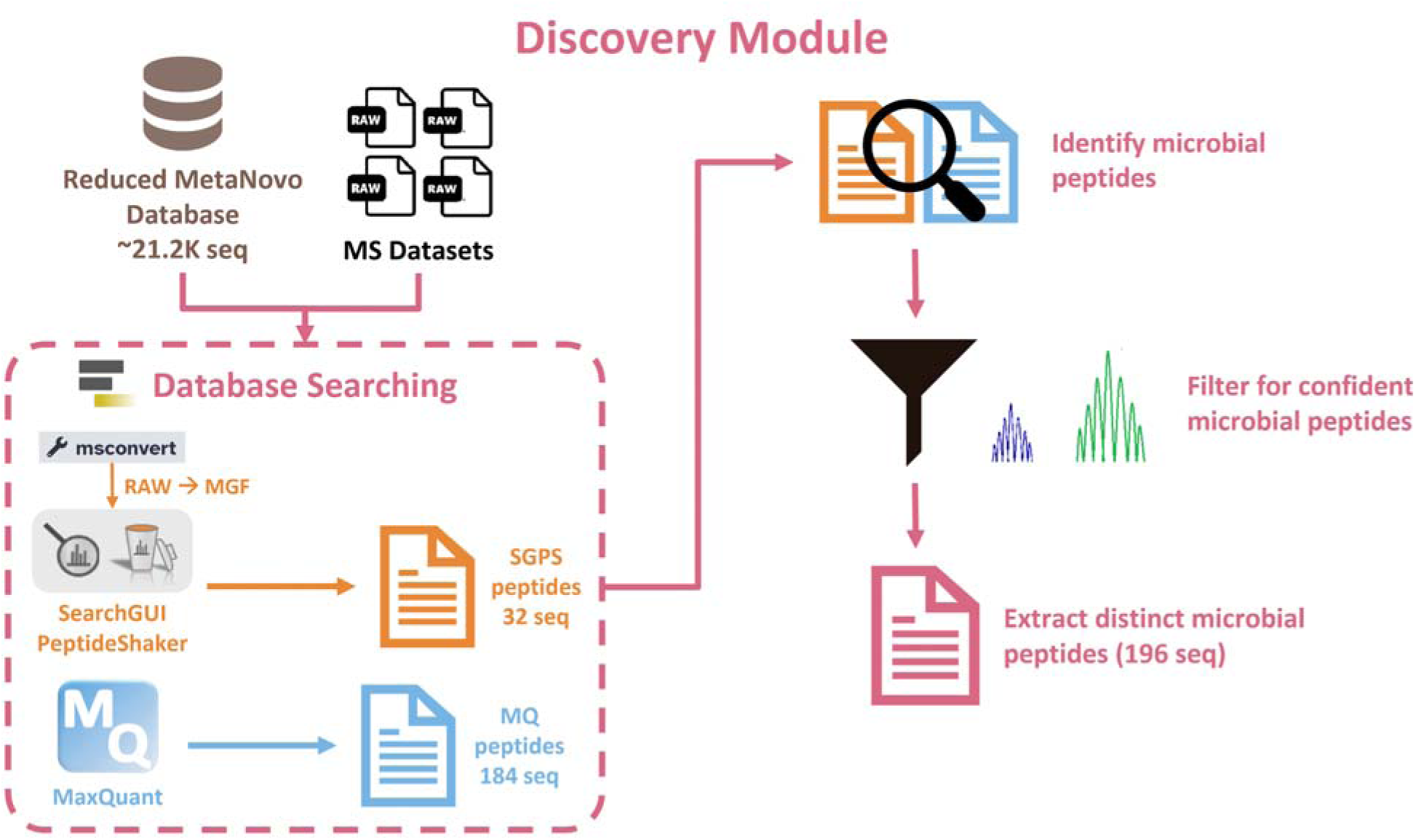
Discovery Module. Microbial peptide identification using SearchGUI/PeptideShaker and MaxQuant.

SearchGUI is a database-searching tool that comprises different search engines to match sample MS/MS spectra to known peptide sequences (14). In this analysis, the search algorithms MS-GF+ and X!Tandem within SearchGUI were employed to match spectra from MS data against peptides from the compact database (15, 16). Subsequently, PeptideShaker was used to organize the detected peptide spectral matches (PSMs), assess the confidence of the data by using false discovery rate (FDR) analysis, and infer the identities of proteins based on the matched peptide sequences (14). Moreover, PeptideShaker generates outputs that can be used to visualize and interpret the data.

MaxQuant is an MS-based proteomics platform that is capable of processing raw data and provides improved mass precision and high precursor mass accuracy, which results in increased protein identification and more in-depth proteomic analysis (17, 18). Following database searching, microbial peptides from SearchGUI/PeptideShaker and MaxQuant were identified, merged, and filtered to retain confident peptides, and grouped to obtain a list of distinct microbial peptides (**Fig. 3**). This list of distinct peptides was then extracted to use as input for PepQuery2 verification.

### Verification module of distinct microbial peptides using PepQuery2

Inputs for the PepQuery2 tool consisted of the list of distinct microbial peptides (from the Discovery module), the four example MS datasets (MGF), the Human UniProt Reference proteome (with isoforms; 82,678 sequences), and contaminants protein sequence databases. PepQuery2 tool further verified the identified microbial peptides detected via the Discovery module to ensure that they were indeed of microbial origin and not a result of human peptides being misassigned (**Fig. 4A**).

**Figure 4.**
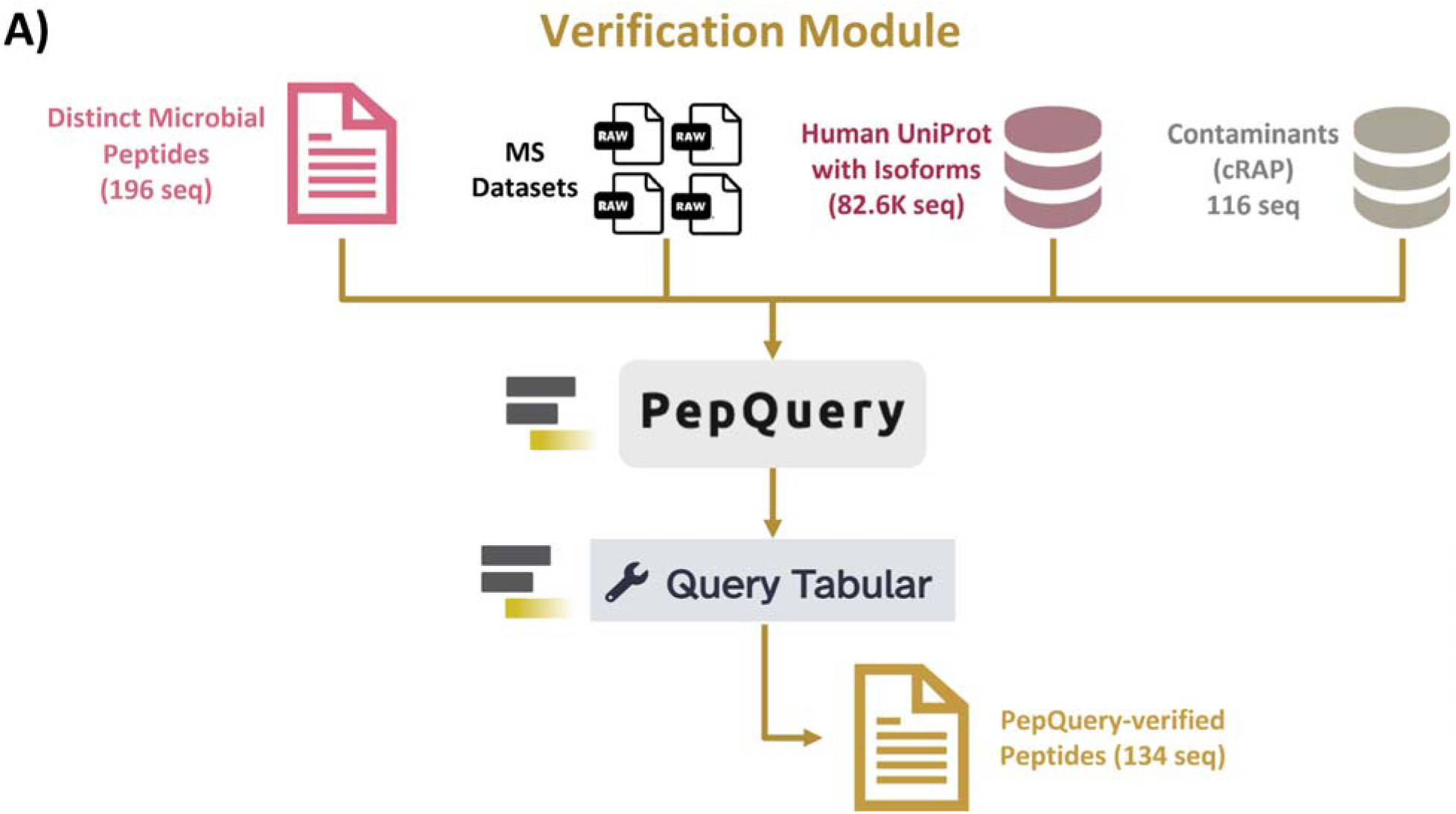
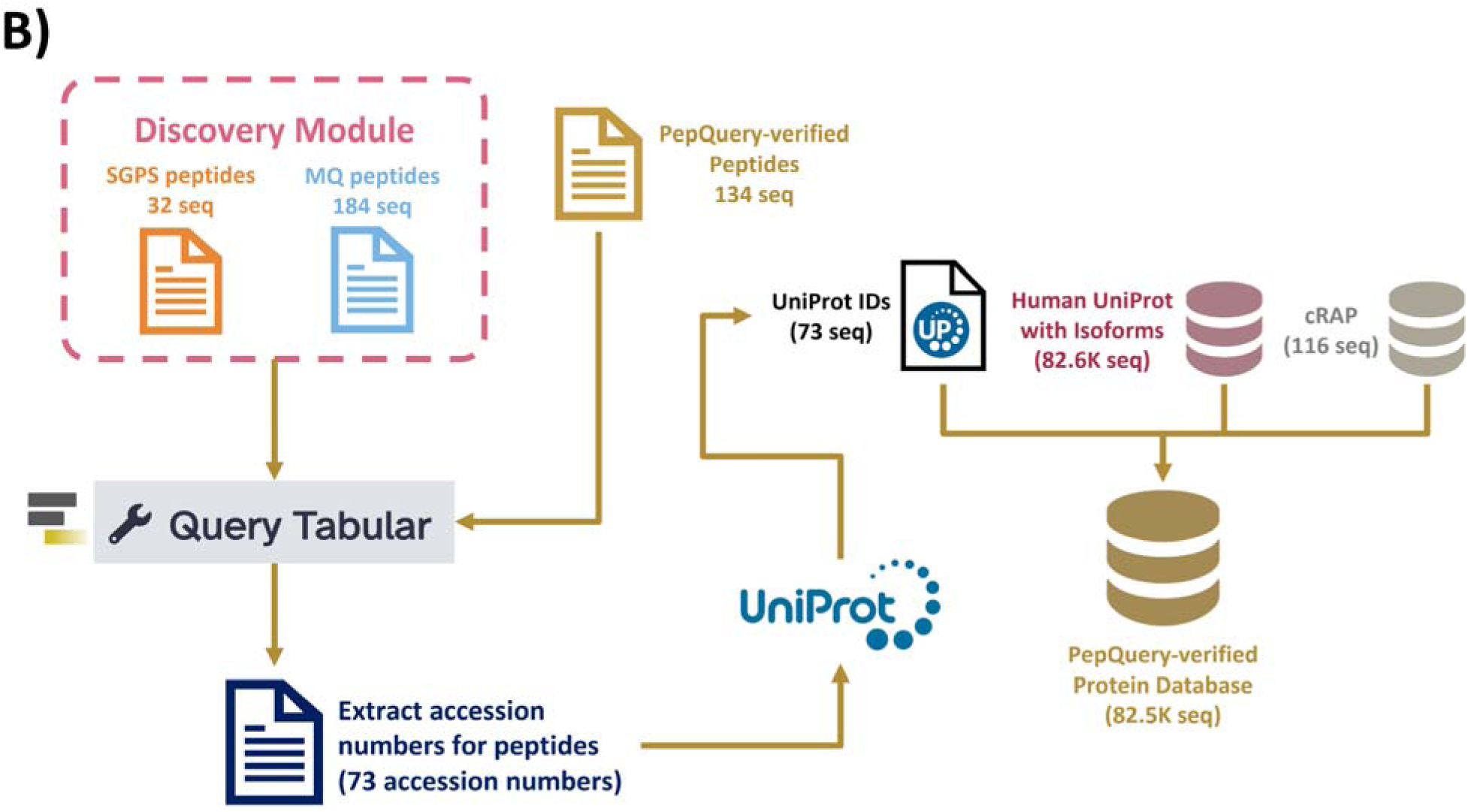
Verification Module. Overview of peptide verification using PepQuery that consists of **A)** generation of PepQuery-verified peptides, which will be used to **B)** generate a PepQuery-verified protein database.

The PepQuery tool was developed as a peptide-centric search engine for MS/MS data analysis to verify the quality of non-host sequences by assigning P-values corresponding to the confidence level in peptide detection (19). PepQuery enables users to search peptide sequences against a proteomics database to verify high–confident PSMs and rigorously examines peptide modifications, which greatly reduces false discoveries in novel peptide identification (19). The PepQuery2 tool builds upon the capabilities of the initial PepQuery software release by providing a new MS/MS spectrum indexing, which results in highly efficient, targeted peptide identification (20). Parameters for PepQuery2 verification in this study are detailed in **Supplementary Table S6**, and both versions of the tool are available as web–based and stand-alone applications (http://www.pepquery.org/).

For each MGF MS dataset used as input, PepQuery2 generated a PSM rank file, each containing peptide identification information. These PSM rank files were compiled and filtered to retain confident PSMs. Then, the Query Tabular tool was used to select microbial peptides detected with confident PSMs and to remove potential contaminants (21). This list of microbial peptides was grouped to obtain distinct peptides (**Fig. 4A**). Then, as shown in **Figure 4B**, PepQuery2-verified peptides and the peptides from SearchGUI/PeptideShaker and MaxQuant (from database searching from the Discovery module) were used as inputs for Query Tabular to extract accession numbers of proteins associated with the PepQuery2-verified peptides. The protein accession numbers of the verified microbial peptides were used to generate a protein sequences FASTA file using the UniProt XML Downloader tool within the Galaxy platform. This microbial protein sequences database (73 sequences) was then merged with the Human UniProt Reference proteome (with isoforms; 82,678 sequences) and cRAP sequence database to generate a protein sequences database (82,562 sequences) for protein and peptide quantification using MaxQuant software (**Fig. 4B**).

### Quantification module using MaxQuant

The quantification module uses the verified microbial database from the previous module along with the Human UniProt and contaminants, RAW MS files, Experimental Design template (for MaxQuant) and Human protein sequences database as inputs (**Fig. 5; Supplementary Table S5**). Proteins and peptides from the MS files were quantified using MaxQuant against the verified microbial protein sequences database (merged with human protein sequences and contaminants sequences; 82,562 protein sequences). This module generated lists of quantified proteins and peptides that were processed for data interpretation and visualization (22). To perform data analysis of the microbial community present in the sample, microbial peptides were extracted from the list of identified peptides. This allowed us to prioritize the identification and quantification of microbial proteins, despite their lower abundance relative to the host proteins in the clinical samples.

**Figure 5.**
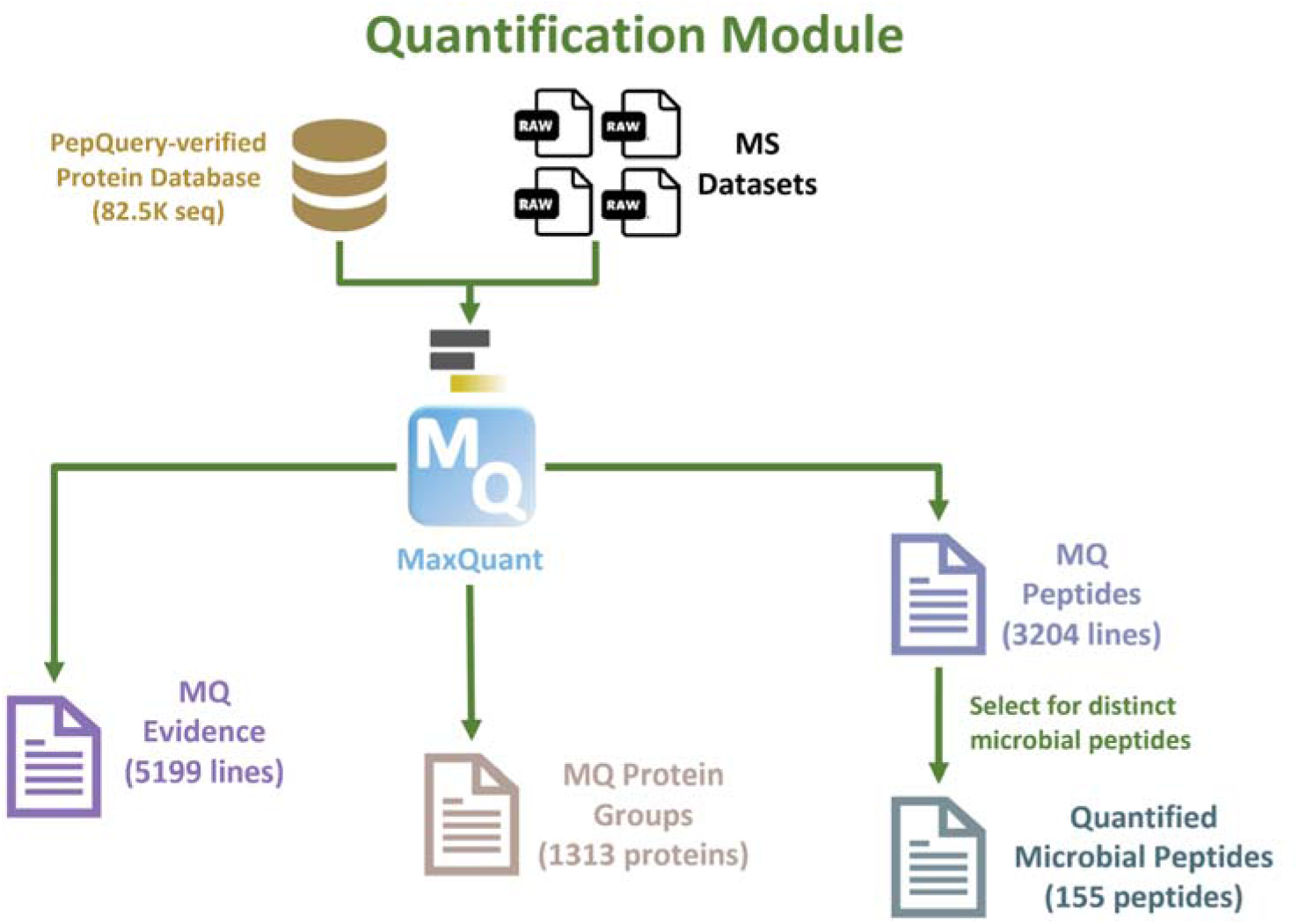
Quantification Module. Overview of peptide quantification using MaxQuant.

### Data interpretation module using Unipept and MSstatsTMT

The protein group text file obtained through MaxQuant quantification was first divided into human and microbial protein groups. To gain deeper insights into the microbial protein groups, the Unipept tool was employed to perform taxonomic and function annotation. First, Unipept determined taxonomic composition by performing lowest common ancestor (LCA) analysis on the peptide sequences, searching them in the UniProt database (7, 23). Unipept annotated the peptides by assigning them to specific taxonomic and functional categories, including the likely genus and species for each microbial peptide sequence (**Fig. 6A**). Unipept outputs included a microbial taxonomic tree and enzyme commission (EC) proteins tree, hierarchical taxonomic annotation with EC numbers, InterPro, and Gene Ontology (GO) terms (**Supplementary Table S7**). These outputs offered a comprehensive understanding of the microbial ecology of the quantified proteins.

**Figure 6.**
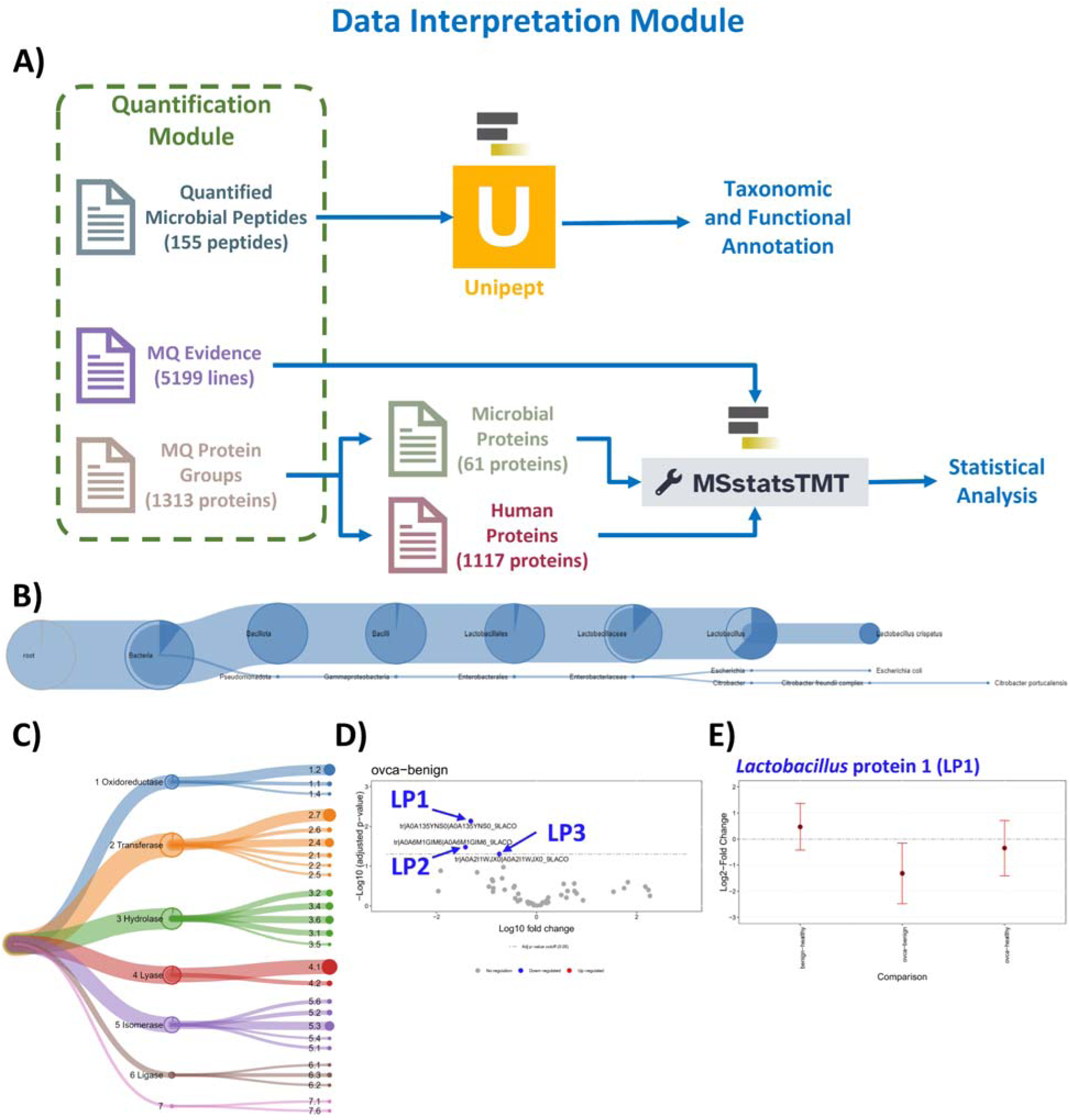
Data Interpretation Module. Overview of data interpretation module using (**A)** Unipept and MSstatsTMT. Examples of the microbial outputs: **(B)** microbial taxonomy tree, (**C**) microbial enzyme commission proteins tree, (**D**) microbial proteins volcano plot (gray denotes no regulation, red/blue for up-/down-regulated), and (**E**) a comparison plot for microbial protein LP1.

Similar to Unipept, the statistical analysis using MSstatsTMT relied on the protein group text file (.txt) obtained through MaxQuant quantification as well as the MaxQuant evidence file (**Fig. 6A; Supplementary Tables S8, S9**). The human and microbial protein groups were separately analyzed using the MSstatsTMT tool. The MSstatsTMT tool is a free, open-source R/Bioconductor package that is compatible with data processing, such as MaxQuant, and allows for sensitive and specific detection of differentially expressed proteins in large-scale experiments with multiple biological samples (22, 24). MSstatsTMT, requiring annotation and comparison matrix files, removed all proteins labeled as contaminants from the MaxQuant protein groups and performed statistical analysis to discern the differential expression of quantified proteins across the three sample groups: healthy, benign, and OC. The annotation file dictated how the quantifications were combined, and the comparison matrix file was needed to accommodate the three different sample groups (OC, healthy, benign). MSstatsTMT generated tabular files with protein abundance ratios, group comparisons with log2 fold changes and P-values, and volcano plots, all of which showcased differentially expressed proteins between human and microbial protein groups. Tutorial material for using MaxQuant and MSstatTMT for TMT data analysis are on the Galaxy Training Network (https://gxy.io/GTN:T00220).

### Data Availability

All input files, including example raw datasets and protein sequence databases, can be accessed on Zenodo via the link: https://doi.org/10.5281/zenodo.10105821. Use the DOI 10.5281/zenodo.10105820 to access the most current version. All information for each module presented in this study is mentioned in the <u>Supplementary Information</u>, including example files and tools (and parameters) hosted on Galaxy Europe Server as well as Galaxy Training Network tutorials deposited on GitHub. All data and links are current as of December 2023.

## RESULTS

### Database generation module: MetaNovo

All modules described in this workflow are summarized in **Supplementary Table S1**, including inputs, software tools, and outputs. The first module of the clinical metaproteomics workflow is the databas generation step, wherein a large database comprising 3,383,217 protein sequences (generated from 118 species) was used (**Fig. 2**). This database was generated using the UniProt XML Downloader tool. This Galaxy tool can also generate a database from proteomes collected at the Genus, Family, Order, or any other higher taxonomy clade. In order to generate a compact protein sequences database, the MetaNovo tool was used. As an input, four example MGF files were processed using the DirecTag tool within MetaNovo to generate a reduced database of 1,908 protein sequences. This reduced database was merged with the Human SwissProt database and contaminants database to generate a database with 21,289 protein sequences so that it could be used for the Discovery module.

### Discovery module: SearchGUI/PeptideShaker and MaxQuant

The discovery module uses the four example RAW files, an experimental design file (for MaxQuant), and MetaNovo protein sequences database (generated from the database generation module) (**Fig. 3**). Using the msconvert tool, MS/MS data (RAW files) were converted to MGF files to search against the MetaNovo-generated database using SearchGUI/PeptideShaker tool. The peptide outputs generated from this module were used for the subsequent verification module. For our example dataset, 184 microbial peptides were detected using MaxQuant, and 32 microbial peptides were detected using SearchGUI/PeptideShaker. Peptides from both tools were merged, filtered, and grouped to retain distinct peptides, and as a result, 196 microbial peptides were identified.

### Verification module: PepQuery2

For verification, 196 microbial peptides (identified in the Discovery module), MGF files, and peptide reports generated from MaxQuant and SearchGUI/PeptideShaker were used as inputs. The module uses the PepQuery2 tool to evaluate MS/MS evidence for microbial sequences identified in the Discovery module. The Query Tabular tool extracted 73 accession numbers of proteins associated with the 134 microbial peptides that passed verification criteria for PepQuery2 (**Fig. 4A**). The 73 accession numbers associated with these verified microbial peptides were used to generate a verified microbial protein sequences database for the Quantification module (**Fig. 4B**).

### Quantification module: MaxQuant

Using the microbial protein sequences containing the verified peptides from PepQuery2, a “compact” database was constructed by merging these with human protein sequences and known contaminants. The MS datasets were searched against this compact database using MaxQuant to generate a peptide report file (3,203 peptides), which was filtered to retain distinct microbial peptides, resulting in a total of 155 quantified microbial peptides (**Fig. 5**). MaxQuant also identified a total of 1,313 protein groups, which included 1,178 non-contaminant protein groups (1,117 human and 61 microbial; **Fig. 5, 6A**).

### Data Interpretation module: Unipept and MSstatsTMT

Unipept provided taxonomic and functional annotation for the 155 quantified microbial peptides (**Fig. 6A**). The taxonomy tree depicted the likely taxonomic assignments for each protein sequence (**Fig. 6B**). *Lactobacillus* was the most abundant genus detected with 90 sequences assigned (56 sequences assigned at genus level and 34 sequences assigned to *L. crispatus*), followed by *Citrobacter* (one sequence assigned to *C. portucalensis*) and *Escherichia* (one sequence assigned to *E. coli*). Unipept also generated an enzyme commission (EC) tree that displays the numerical classification of what chemical reaction is catalyzed. The most prominent classification was transferase with 33 sequences, followed by hydrolase with 29 sequences and lyase with 24 sequences (**Fig. 6C**).

The protein group text file obtained through MaxQuant quantification contained 1,313 protein groups. After selecting human and microbial protein groups, this resulted in 1,117 human and 61 microbial proteins (**Fig. 6A**). Additionally, MSstatsTMT generated volcano plots (**Fig. 6D**) and comparisons (OC, healthy, benign) to identify differentially expressed microbial proteins (**Fig. 6E, Supplementary Fig. S1– S2**). As seen in **Figure 6D**, a volcano plot visualizes the log10-fold changes and P-values for a comparison between OC and benign cases for the 62 microbial proteins. The horizontal dashed line represents the FDR cutoff (P–value = 0.05), and data points above this line denote statistically significant proteins. The data points are colored to denote any differential expression between cases: gray for no regulation and red/blue for up-/down-regulation. In this study, three microbial proteins from *Lactobacillus* were determined to be statistically significant when comparing OC versus benign cases. For demonstration purposes, these proteins have been labeled *Lactobacillus* protein (LP) 1, LP2, LP3 in **Figure 6D**. As seen in **Figure 6E**, a comparison plot visualizes log2-fold changes and variation of multiple comparisons for a single protein. An example plot for one microbial protein that was statistically significant when comparing the OC cases to the benign cases was LP1 (**Fig. 6E**).

## DISCUSSION

Mass spectrometry-based clinical metaproteomics, which offers insights into the functional molecules expressed by microorganisms as well as taxonomic composition, within clinical samples has been gaining attention in recent years (1, 25, 26). Moreover, the approach also offers insights into how the host environment responds to microbial infection, via simultaneous analysis of the host proteome. Clinical metaproteomics has been used in unraveling pathogenic mechanisms in Alzheimer’s disease, autism, colorectal cancer, cystic fibrosis, diabetes, inflammatory bowel disease, and COVID-19 co-infection in a variety of clinical sample types including: feces, bronchoalveolar lavage fluid, vaginal swabs, and oral cavity specimens (27–40).

Despite its immense potential in offering functional insights into the microbiome as well as host response to infection, the clinical metaproteomics field faces some challenges. These include challenges in experimental design, protein sequence database composition, computational tools and resources for raw data processing, functional and taxonomy annotation, data variability, and training resources for non-experts (1). Foremost amongst these challenges is the relatively low abundance of microbial proteins as compared to the host proteins in clinical samples, thus making it difficult to detect and characterize these microbial proteins (27, 31). While developing improved analytical or instrumental methods to enrich low–abundance microbial proteins or to enhance sensitivity to detect these proteins in complex clinical samples is beyond the scope of this study, we have developed a bioinformatics workflow that enables improved characterization of microbial proteins that can be detected in clinical samples. Additionally, this bioinformatics workflow is not limited to only microbial proteins as it can be adapted for analysis of human proteins (**Supplementary Fig. S3**).

As a first step, we have addressed the issue of large protein sequence databases that are used in metaproteomics studies. While there are multiple database generation methods, including iterative approaches, we used the MetaNovo approach to generate a reduced database for searching (41–44). MetaNovo uses *de novo* sequence tag matching along with an algorithm for probabilistic optimization of the large input database to generate a customized protein sequence database. In order to further improve the chances of detection of microbial proteins, we have used multiple database search algorithms for detection of confident PSMs. While SearchGUI/PeptideShaker offers multiple search algorithms, we have used X!Tandem and MS-GF+ for demonstration purposes in this study. Users can also try other search algorithms such as Comet within SearchGUI/PeptideShaker. We have also used MaxQuant within Galaxy to search for additional microbial peptides from the same sample. In the future, search algorithms such as FragPipe (45) and Scribe (46) will also be available within the Galaxy framework and add to the repertoire of tools and workflows for advanced multiomics analysis (9).

One step that is often overlooked when using MS-based metaproteomics is the need to ascertain the quality of the PSMs for microbial peptide identification and protein inference. This is especially important since most of the modern search algorithms rely on FDR analysis for the identification of peptides and proteins from data-dependent-acquisition (DDA) MS data (47). This might lead to false positive PSMs, especially from microbes of low abundance that are repeatedly identified using large protein sequence databases, which can erroneously bolster the number and quality of PSMs. Therefore, there is a need to verify these proteins using bioinformatic approaches which can lead to further validation via targeted proteomic methods (48, 49). Various methods such as Peptide-Spectral-Match-Evaluation and those within the Multi-omics Visualization Platform are available for evaluation within the Galaxy platform (50, 51). For this workflow, we have used the PepQuery2 tool which verifies novel and known peptides identified using spectrum-centric database searching (20). The verification module helps to further reduce the database size, enabling quantitative analysis.

For the quantification module, we have used MaxQuant software within Galaxy which generates quantitative protein and peptide outputs ready for MSstatsTMT analysis in the data interpretation module (22). Although MaxQuant was used in this study, newer software such as FragPipe and Scribe are planned for Galaxy implementation and could also be used for quantitative analysis (45, 46). The data interpretation module uses MSstatsTMT and Unipept which enable data interpretation via visualization (23, 52). We have described the use of variations of this interpretation module for prior clinical metaproteomics studies (10).

The workflow we describe here is built with an eye towards flexibility to incorporate emerging methods and technologies that may further improve clinical metaproteomic studies. We believe that the outputs from the modules described above will aid in prioritizing proteins and peptides of highest interest. The workflow outputs can be used to provide the necessary information needed to develop targeted assays for validation and possible implementation within the clinic. For example, methods for enriching microbial proteins from clinical samples containing high–abundance host proteins would enhance depth of detection via MS as well as provide improved starting data, immediately compatible with our workflow. Metaproteomics researchers have also started using data-independent acquisition for quantitative studies, with growing evidence indicating that this approach can greatly improve the depth and quantitative accuracy of measurements (37, 53, 54). Along with the sensitive mass spectrometers available now for deep quantification studies (37, 55), this offers an opportunity for metaproteomics researchers to go deeper into the microbiome functions along with deciphering taxonomic contributions.

To efficiently process and analyze large-scale datasets, metaproteomic researchers must employ software and hardware that are capable of scalability to accommodate memory- and compute-intensive tools and resources for raw data processing, especially with more complex workflows (50, 56). We addressed this challenge with the Galaxy implementation for the development of this workflow as Galaxy provides extensive access to tools and data that can be implemented with minimal training (8, 10, 50). The Galaxy ToolShed operates as a massive tool repository that can be used by developers and admins to host, share, and install thousands of free Galaxy tools into their instances (>9,000 tools, as of Sept. 2023; https://toolshed.g2.bx.psu.edu/). The ToolShed facilitates reproducible analyses by tracking software versions (57). Coupled with the flexible, modular nature of Galaxy, users are able to easily select, test, and replace tools as new tools are made available to best suit their analyses. The Galaxy platform is accessible via high–performance computing (HPC) systems and public cloud environments (22).

Possessing the computational tools to process large–scale datasets does not automatically mean users are immediately prepared to perform their own data analyses if they cannot understand what they are doing. Comprehensive training, such as basic computational skills and data analysis, remains a requirement for understanding data analyses and result interpretation (6). However, even as the life sciences are continually evolving to be more computational in nature and bioinformatics becoming more prevalent in research studies, life science educational programs do not commonly include comprehensive computational training in their curricula (6). To address this training deficit, the GTN (https://training.galaxyproject.org/) was created to provide learners and instructors, regardless of expertise and experience, with free online training materials and access to a global community, while promoting open data analysis practices (6). Training materials, covering topics from the life sciences to machine learning, include tutorials, video lecture recordings, and coding tutorials, as well as Train-the-trainer (TtT) tutorials for instructors and workshop resources (6).

Due to its wide array of tools and user-friendly interface, the Galaxy platform holds great potential for researchers as well as for students and other non–experts. Similarly to how bioinformatics has become more prominent in research studies, education has increasingly shifted to online formats, most notably with the surge of converting to online learning due to the COVID–19 pandemic (6, 58). The Galaxy platform’s flexibility accommodates self–paced and explorative learning and combats learner isolation with its global community of fellow learners, contributors, and instructors (58). The Galaxy platform empowers learners and researchers worldwide, regardless of expertise, with the tools and skills to perform their own data analyzes, all readily accessible through a standard web browser.

In summary, we have developed a bioinformatics clinical metaproteomics workflow within the Galaxy platform. This workflow will enable metaproteomics researchers to generate customized search databases, search MS data using multiple search algorithms, verify the quality of spectral matches, quantify proteins and peptides, and interpret the data using generated visualizations. We anticipate that the availability of this workflow will enable metaproteomics researchers to undertake challenging clinical metaproteomics investigations.

## Supporting information

Clinical MP Supplemental

## ACKNOWLEDGEMENTS

We thank Dr. Kristin Boylan at the University of Minnesota and members of the Pacific Northwest National Laboratories, especially Drs. Paul Piehowski, Tao Liu, and Karin Rodland for their technical expertise in the collection and processing of the Pap test samples and generation of the TMT MS data used in this study. This project was funded in part by the Minnesota Ovarian Cancer Alliance, the National Institutes of Health/National Cancer Institute Grant Number: 5R01CA262153 (A.P.N.S.), and The National Institutes of Health/National Cancer Institute Grant Number: P30CA077598 (P.D.J. and T.J.G.).

## SUPPLEMENTARY INFORMATION

- Clinical MP Manuscript Supplemental

