## Supplementary material for "A novel clinical metaproteomics workflow enables bioinformatic analysis of host-microbe dynamics in disease": Clinical MP Supplemental

**Supplementary Information**

**Section 1. Galaxy Materials**

**Table S1. Module Summary: Inputs, Software Tools, and Outputs.** This table compiles inputs, outputs, and software tools for each module.

| **MODULE** | **INPUTS** | **SOFTWARE TOOLS** | **OUTPUTS** |
| --- | --- | --- | --- |
| **DATABASE GENERATION** | 4 MGF Files.  Human SwissProt Database (reviewed-only; 20,408 protein sequences, as of Sept. 2023).  Contaminant protein sequences (116 protein sequences).  Protein sequences database of 118 species (3,383,217 protein sequences). | FASTA Merge Files and Filter Unique Sequences.  MetaNovo. | MetaNovo generated a protein sequences database for Discovery Module (21,289 protein sequences) |
| **DISCOVERY** | 4 RAW Files.  Human SwissProt Database (reviewed-only; 20,408 protein sequences, as of Sept. 2023).  Contaminant protein sequences (116 protein sequences).  Experimental Design Template.  MetaNovo generated a protein sequences database for Discovery Module (21,289 protein sequences). | msconvert.  Protein Database Downloader.  FASTA Merge Files and Filter Unique Sequences.  FASTA To Tabular Converter.  Filter Tabular.  FastaCLI.  Identification Parameters.  SearchGUI.  PeptideShaker.  MaxQuant.  Select lines that match an expression.  Filter data on any column using simple expressions.  Query Tabular using sqlite SQL.  Cut.  Remove Beginning.  Concatenate datasets.  Group data by a column and perform aggregate operation on other columns. | Distinct microbial peptides for PepQuery2 (196 peptide sequences) |
| **VERIFICATION** | 4 MGF Files.  Distinct microbial peptides for PepQuery2 (196 peptide sequences).  PeptideShaker Peptide Report (1,428 lines)  MaxQuant Peptide Report (3,284 lines).  Human UniProt (+Isoforms) (82,678 sequences).  Contaminant protein sequences (116 protein sequences). | Protein Database Downloader.  FASTA Merge Files and Filter Unique Sequences.  PepQuery2.  Collapse Collection into single dataset in order of the collection.  Filter data on any column using simple expressions.  Query Tabular using sqlite SQL.  Cut.  Remove Beginning.  Concatenate datasets.  Group data by a column and perform aggregate operation on other columns.  UniProt download proteome as XML or fasta. | Protein sequences database for Quantification Module (82,562 protein sequences). |
| **QUANTIFICATION** | 4 RAW Files.  Experimental Design Template.  Protein sequences database for Quantification Module (82,562 protein sequences). | MaxQuant.  Select lines that match an expression.  Cut columns from a table.  Group data by a column and perform aggregate operation on other columns. | Microbial peptides with quantification (155 peptide sequences)  Microbial proteins with quantification (61 protein sequences)  MaxQuant Protein Groups (1,313 protein sequences).  MaxQuant Evidence (5,199 lines). |
| **DATA INTERPRETATION** | Microbial peptides with quantification (155 peptide sequences).  MaxQuant Protein Groups (1,313 protein sequences).  MaxQuant Evidence (5,199 lines).  Annotation.  Comparison Matrix. | Select lines that match an expression.  Unipept retrieve taxonomy for peptides.  MSstatsTMT protein significance analysis in shotgun mass spectrometry-based proteomic experiments with tandem mass tag (TMT) labeling. | 61 microbial proteins  1,117 human proteins  Microbial Taxonomy Tree.  Microbial EC Proteins Tree.  Microbial Proteins Volcano Plot.  Microbial Proteins Comparison. |

**Table S2. Galaxy History Links: Inputs, Tools, Outputs.** This table summarizes the Galaxy history links for the five modules, including inputs, workflows, and outputs. Note that link contents, such as number of sequences in databases, tool versions, are current at the time of writing (December 2023) but may vary as Galaxy and associated tools and databases are updated.

|  | **INPUT** | **TOOLS** | **OUTPUT** |
| --- | --- | --- | --- |
| **DATABASE GENERATION** | <https://usegalaxy.eu/u/galaxyp/h/input1databasegeneration> | <https://usegalaxy.eu/u/galaxyp/w/wf1databasegenerationworkflow> | <https://usegalaxy.eu/u/galaxyp/h/outputwf1databasegeneration> |
| **DISCOVERY** | <https://usegalaxy.eu/u/galaxyp/h/input2discovery-1> | <https://usegalaxy.eu/u/galaxyp/w/wf2discovery-workflow> | <https://usegalaxy.eu/u/galaxyp/h/wf2outputdiscovery> |
| **VERIFICATION** | <https://usegalaxy.eu/u/galaxyp/h/input3verification> | <https://usegalaxy.eu/u/galaxyp/w/wf3verificationworkflow> | <https://usegalaxy.eu/u/galaxyp/h/wf3outputverification> |
| **QUANTIFICATION** | <https://usegalaxy.eu/u/galaxyp/h/input4quantitation-1> | <https://usegalaxy.eu/u/galaxyp/w/wf4quantitationworkflow> | <https://usegalaxy.eu/u/galaxyp/h/wf4outputquantitation> |
| **DATA INTERPRETATION** | <https://usegalaxy.eu/u/galaxyp/h/input5datainterpretation-1> | <https://usegalaxy.eu/u/galaxyp/w/wf5datainterpretationworklow> | <https://usegalaxy.eu/u/galaxyp/h/wf5outputdatainterpretation> |

**Table S3. Galaxy Training Network (GTN) materials.** This table contains the GTN links where users will be able to access and perform the analysis as instructed. Note that links are up to date at time of writing (Dec. 2023).

| **ALL INPUT FILES** | <https://zenodo.org/records/10105821> |
| --- | --- |
| **DATABASE GENERATION** | <https://github.com/subinamehta/training-material/blob/main/topics/proteomics/tutorials/clinical-mp-database-generation/tutorial.md> |
| **DISCOVERY** | <https://github.com/subinamehta/training-material/blob/main/topics/proteomics/tutorials/clinical-mp-discovery/tutorial.md> |
| **VERIFICATION** | <https://github.com/subinamehta/training-material/blob/main/topics/proteomics/tutorials/clinical-mp-data-verification/tutorial.md> |
| **QUANTIFICATION** | <https://github.com/subinamehta/training-material/blob/main/topics/proteomics/tutorials/clinical-mp-quantitation/tutorial.md> |
| **DATA INTERPRETATION** | <https://github.com/subinamehta/training-material/blob/main/topics/proteomics/tutorials/clinical-mp-data-interpretation/tutorial.md> |

**Section 2. Discovery Search Engine Parameters**

**Table S4. SearchGUI/PeptideShaker and MaxQuant Parameters.**

|  | **SearchGUI/PeptideShaker** | **MaxQuant** |
| --- | --- | --- |
| **Enzyme** | Trypsin | Trypsin/P |
| **Enzyme specificity** | Specific at both termini | Specific |
| **Precursor tolerance (ppm)** | 10.0 | 10.0 |
| **Fragment tolerance (Da)** | 0.6 | 0.6 |
| **Allowed missed cleavages** | 2 | 2 |
| **Minimum peptide length** | 8 | 8 |
| **Maximum peptide length** | 30 | 50 |
| **Modifications** | Fixed: Carbamidomethylation of C, TMT 11-plex of K+4, TMT 11-plex of peptide N-term  Variable: Oxidation of M | Fixed: Carbamidomethyl (C)  Variable: Oxidation (M) |
| **Decoy parameters** | Decoy flag: _REVERSED,  Target decoy suffix: _concatenated_target_decoy | Decoy mode: Revert |
| **False Discovery Rate (FDR)** | Protein FDR: 1.0%  Peptide FDR: 1.0%  PSM FDR: 1.0% | Protein FDR: 0.01 (1.0%)  PSM FDR: 0.01 (1.0%) |
| **Other:** | Search engines: X!Tandem, MS-GF+ | Isobaric Labeling: tmt11plex |

**Table S5. MaxQuant Experimental Design Template For Discovery and Quantification.**

| **Name** | **Fraction** | **Experiment** | **PTM** |
| --- | --- | --- | --- |
| PTRC_Skubitz_Plex2_F10_9Aug19_Rage_Rep-19-06-08.raw | 1 | 1 | FALSE |
| PTRC_Skubitz_Plex2_F11_9Aug19_Rage_Rep-19-06-08.raw | 2 | 1 | FALSE |
| PTRC_Skubitz_Plex2_F13_9Aug19_Rage_Rep-19-06-08.raw | 3 | 1 | FALSE |
| PTRC_Skubitz_Plex2_F15_9Aug19_Rage_Rep-19-06-08.raw | 4 | 1 | FALSE |

**Section 3. Verification Parameters**

**Table S6. PepQuery2 Parameters.**

| **Parameter** | **Value** |
| --- | --- |
| **Fixed Modifications** | 1: Carbamidomethylation of C [57.02146372057]  13: TMT 11-plex of K [229.16293213472]  14: TMT 11-plex of peptide N-term [229.16293213472] |
| **Variable Modifications** | 2: Oxidation of M [15.99491461956] |
| **Use more stringent criterion for unrestricted modification searching** | Yes |
| **Consider amino acid substitution modifications?** | Yes |
| **Digestion Enzyme** | Trypsin |
| **Max Missed Cleavages** | 2 |
| **Precursor Tolerance (ppm)** | 10 |
| **Fragment Tolerance (Da)** | 0.6 |
| **Fragmentation Method** | CID/HCD |
| **Scoring Method** | HyperScore |
| **Select outputs** | psm_rank.txt |

**Section 4. Data Interpretation Parameters**

**Table S7. Unipept parameters.**

| **Parameter** | **Value** |
| --- | --- |
| **Unipept application** | Peptinfo: Tryptic peptides and associated EC and GO terms and lowest common ancestor taxonomy |
| **Equate isoleucine and leucine**  (isoleucine (I) and leucine (L) are equated when matching tryptic peptides to UniProt records) | Yes |
| **Retrieve extra information**  (Return the name of the EC-number and GO-term) | Yes |
| **Group responses by GO namespace (biological process, molecular function, cellular component)** | No |
| **Names**  (Return the names in complete taxonomic lineage) | Yes |
| **Match input peptides by:** | Match to the full input peptide |
| **Show tryptic_match as:** | List of matched tryptic parts |
| **Outputs** | Tabular with one line per peptides  JSON Taxomony Tree (for pept2lca, pep2taxa, and peptinfo)  Peptide GO terms in normalized tabular (for pept2go, pept2funct, and peptinfo)  Peptide InterPro entries in normalized tabular (for pept2interpro, pept2funct, and peptinfo)  Peptide EC terms in normalized tabular (for pept2ec, pept2funct, and peptinfo)  JSON EC Coverage Tree (for pept2ec, pep2funct, and peptinfo) |

**Table S8. Excerpt from MSstatsTMT annotation file.** This table depicts an excerpt from the MSstatsTMT annotation file for each run (RAW file). The complete file includes the same information for all four runs.

| **Run** | **Fraction** | **TechRepMixture** | **Channel** | **Condition** | **BioReplicate** | **Mixture** |
| --- | --- | --- | --- | --- | --- | --- |
|  | 1 | 1 | channel1 | healthy | 1 | 1 |
|  | 1 | 1 | channel2 | ovca | 1 | 1 |
|  | 1 | 1 | channel3 | benign | 1 | 1 |
|  | 1 | 1 | channel4 | ovca | 1 | 1 |
|  | 1 | 1 | channel5 | healthy | 1 | 1 |
|  | 1 | 1 | channel6 | ovca | 1 | 1 |
|  | 1 | 1 | channel7 | benign | 1 | 1 |
|  | 1 | 1 | channel8 | ovca | 1 | 1 |
|  | 1 | 1 | channel9 | benign | 1 | 1 |
|  | 1 | 1 | channel10 | ovca | 1 | 1 |
|  | 1 | 1 | channel11 | norm | 1 | 1 |

**Table S9. MSstatsTMT Comparison Matrix file.**

| **names** | **ovca** | **healthy** | **benign** | **norm** |
| --- | --- | --- | --- | --- |
| **ovca-healthy** | 1 | -1 | 0 | 1 |
| **ovca-benign** | 1 | 0 | -1 | 1 |
| **benign-healthy** | 0 | -1 | 1 | 1 |
| **norm** | 1 | 1 | 1 | 1 |

**Section 5. Additional Examples of Data Interpretation Outputs**

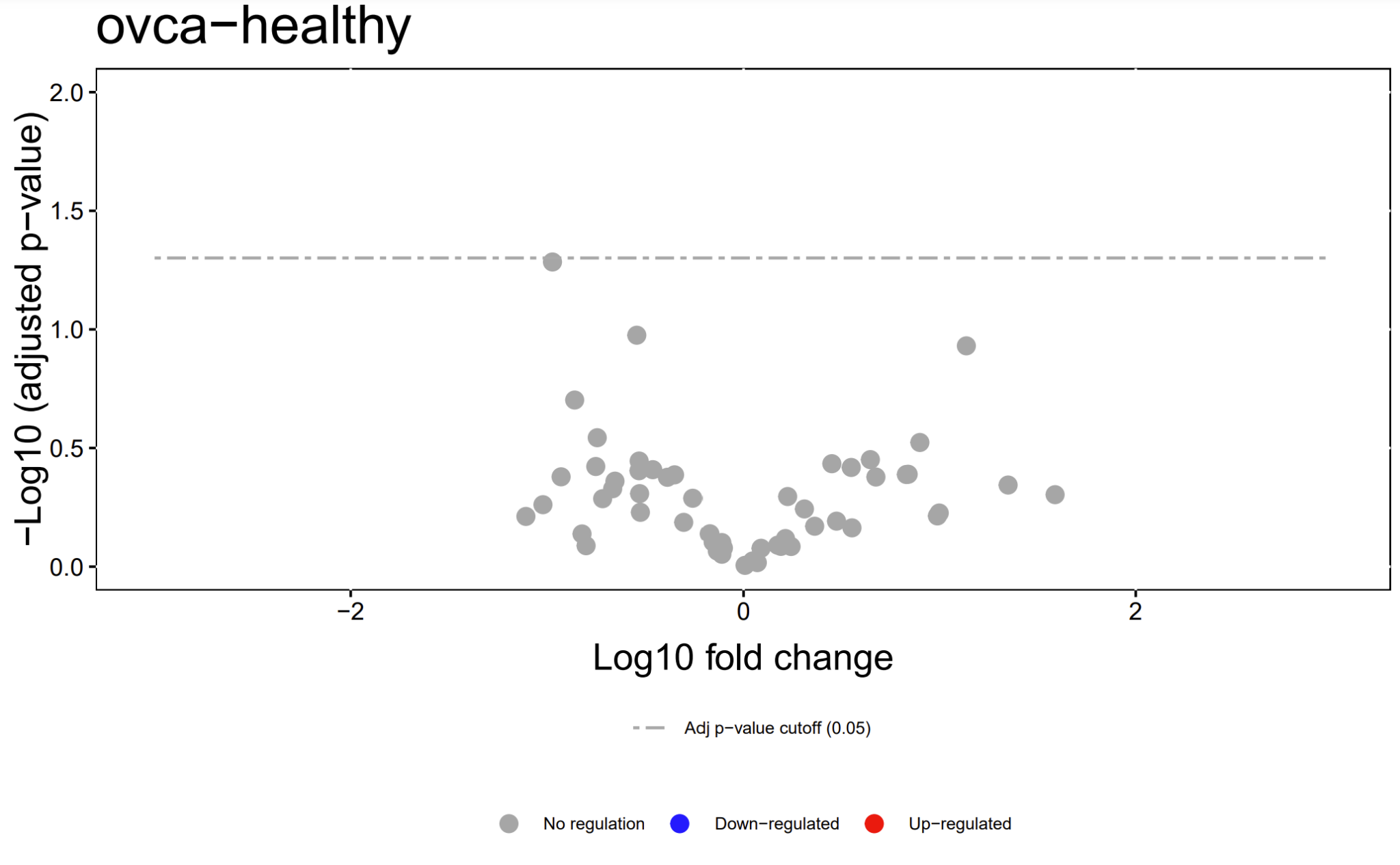

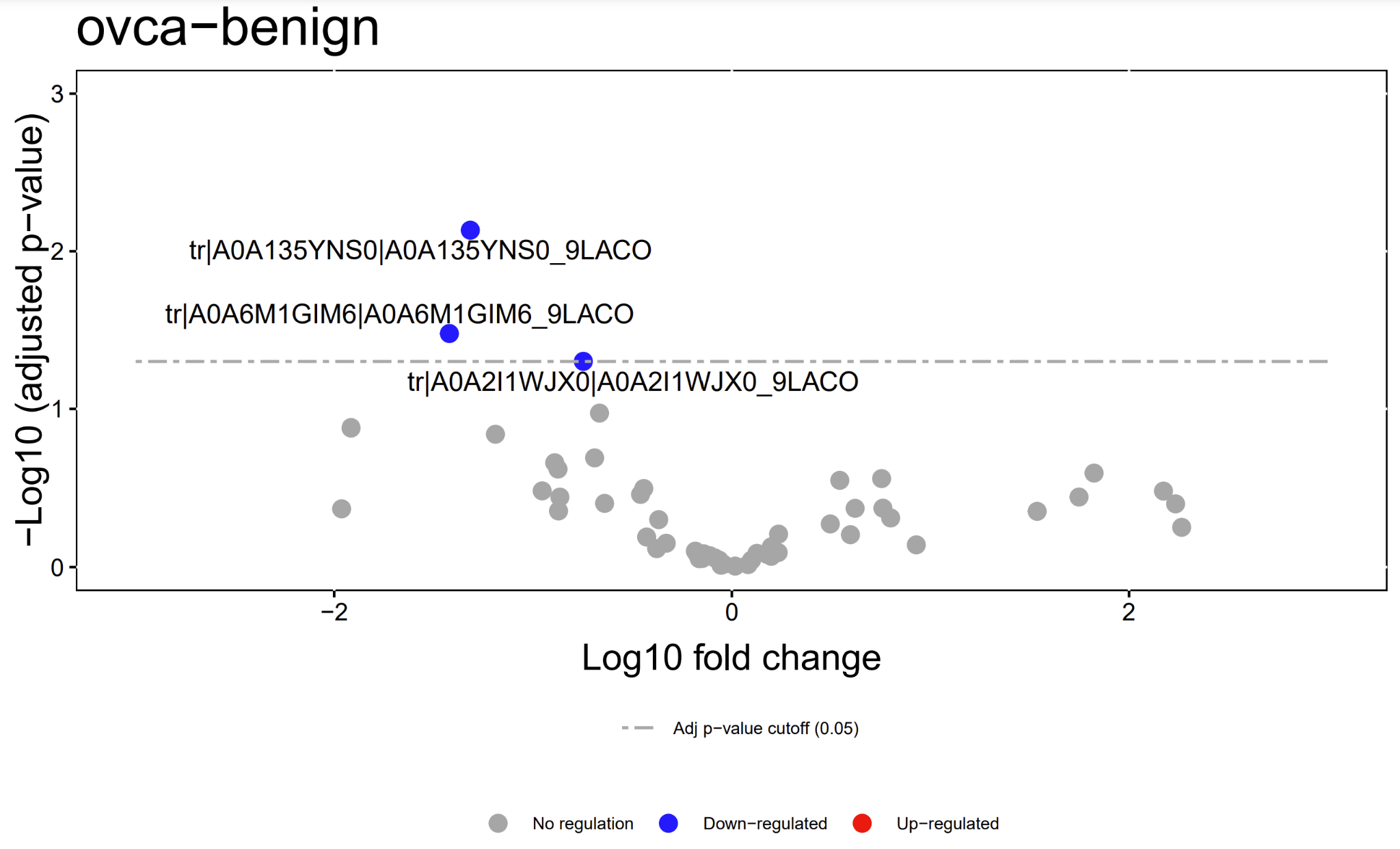

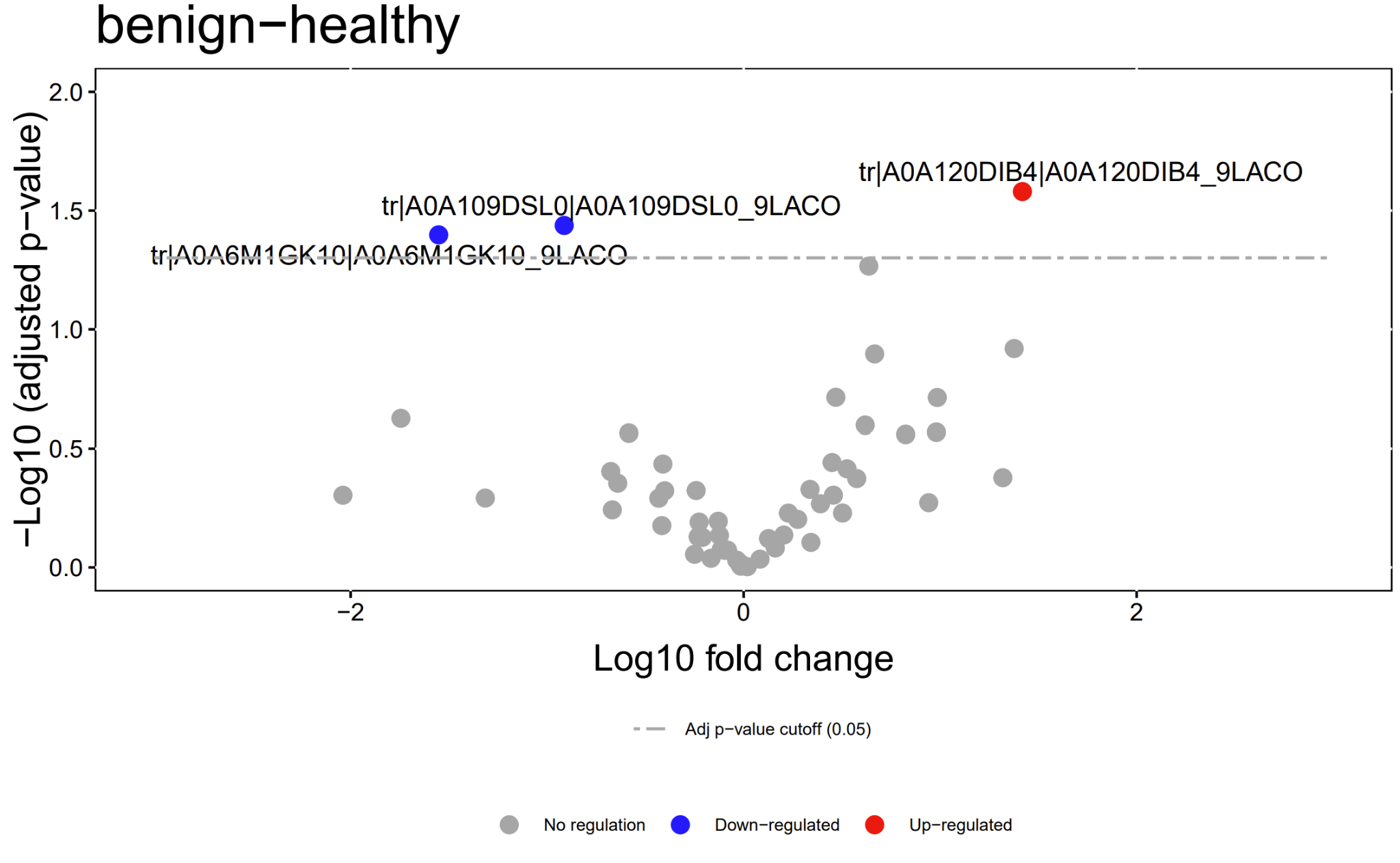

**Figure S1. Microbial Protein Volcano Plots.** Examples of volcano plots illustrating the log10 fold changes and P-values for comparisons (ovca–healthy, ovca–benign, benign–healthy) for quantified microbial proteins.

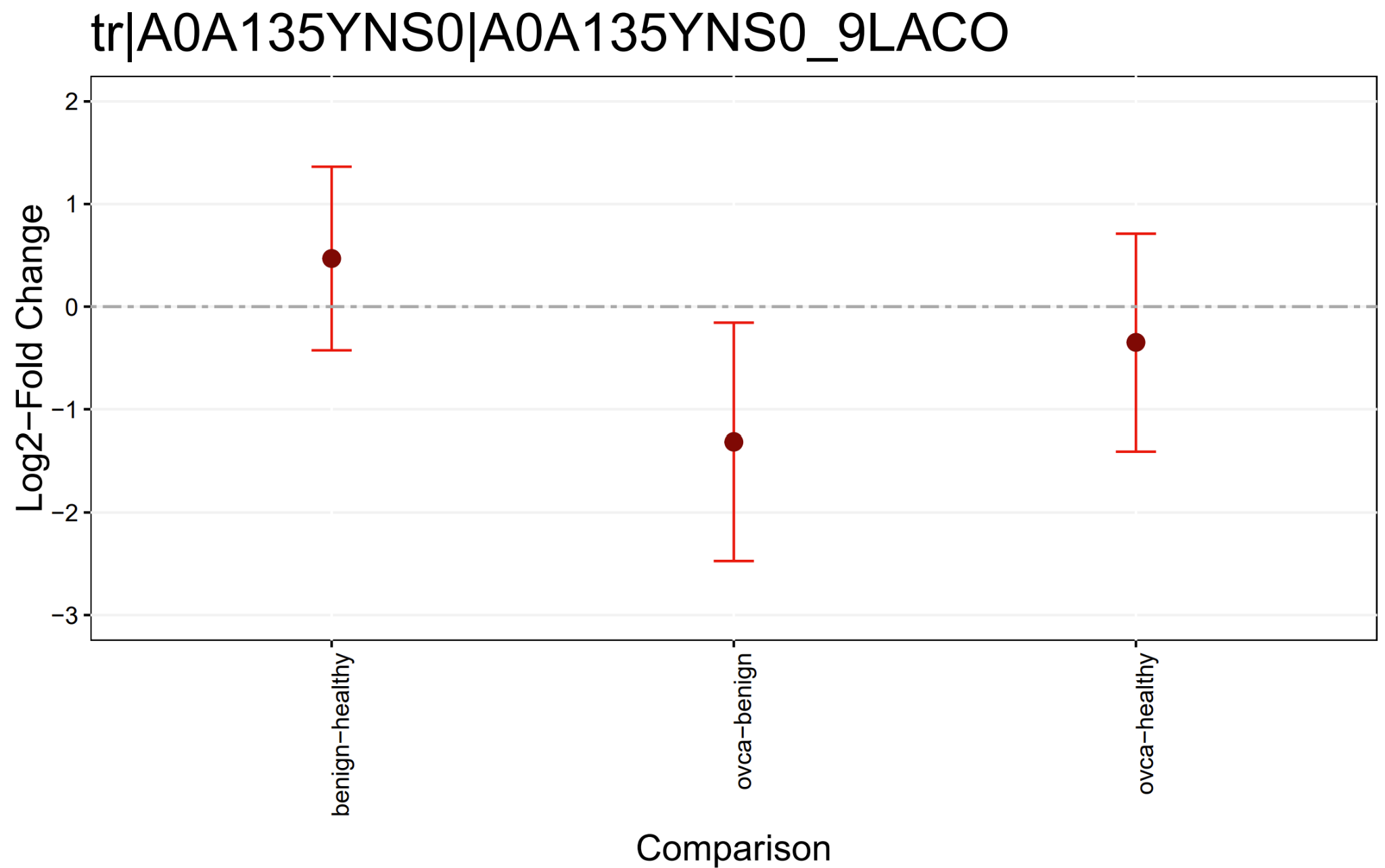

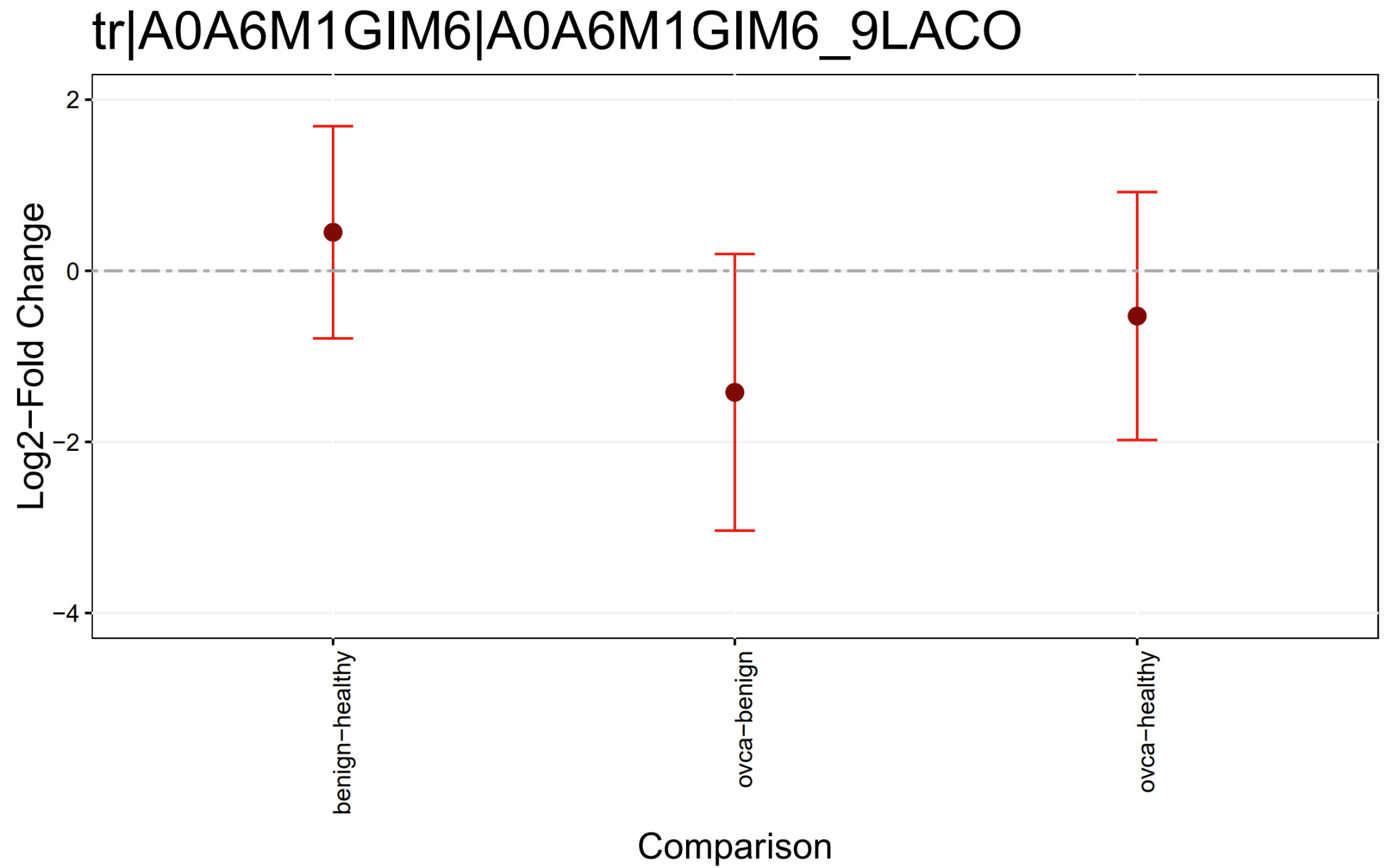

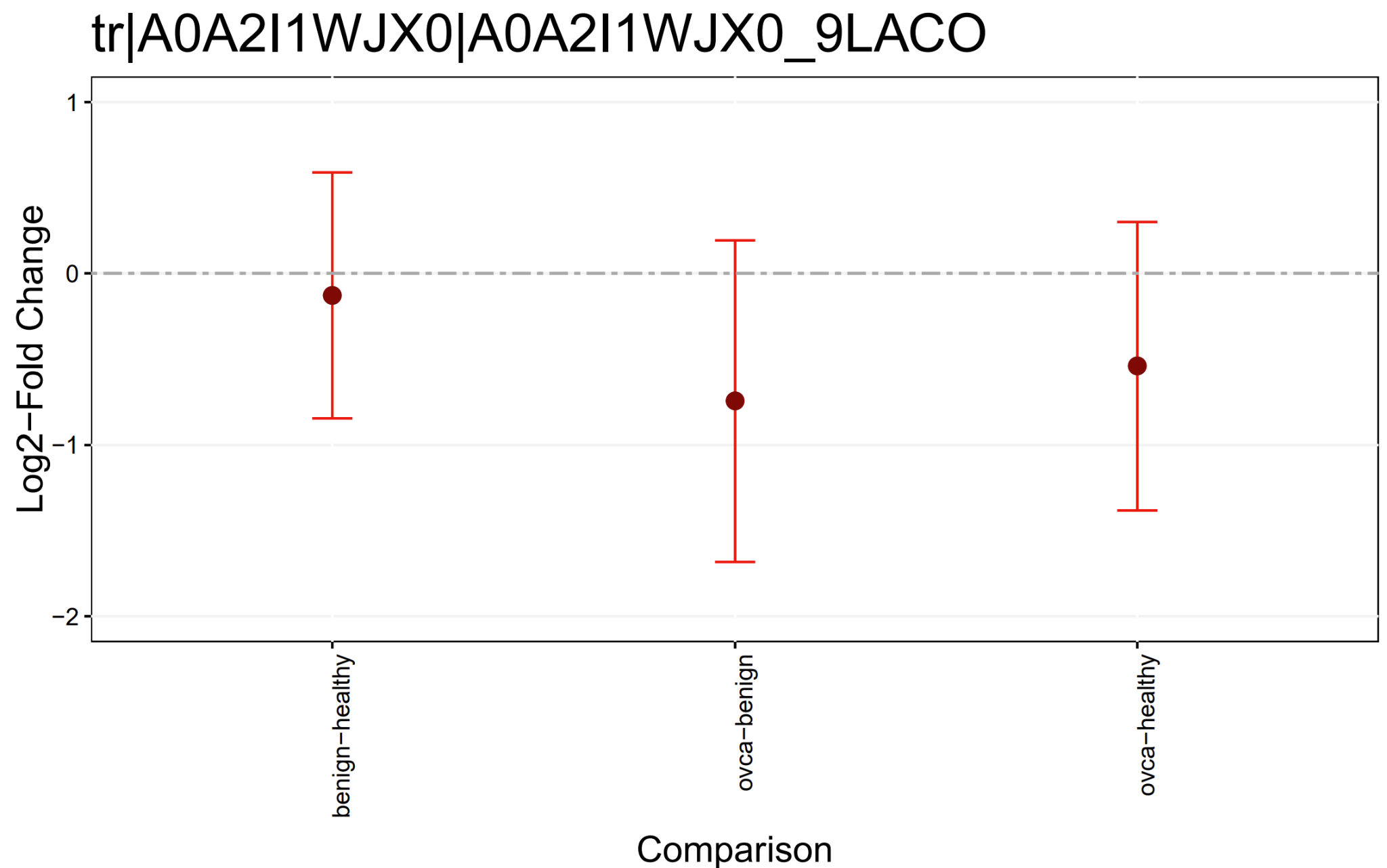

**Figure S2. Microbial Comparison Plots.** Examples of comparison plots visualizing log2 fold changes and variation of multiple comparisons for quantified microbial proteins.

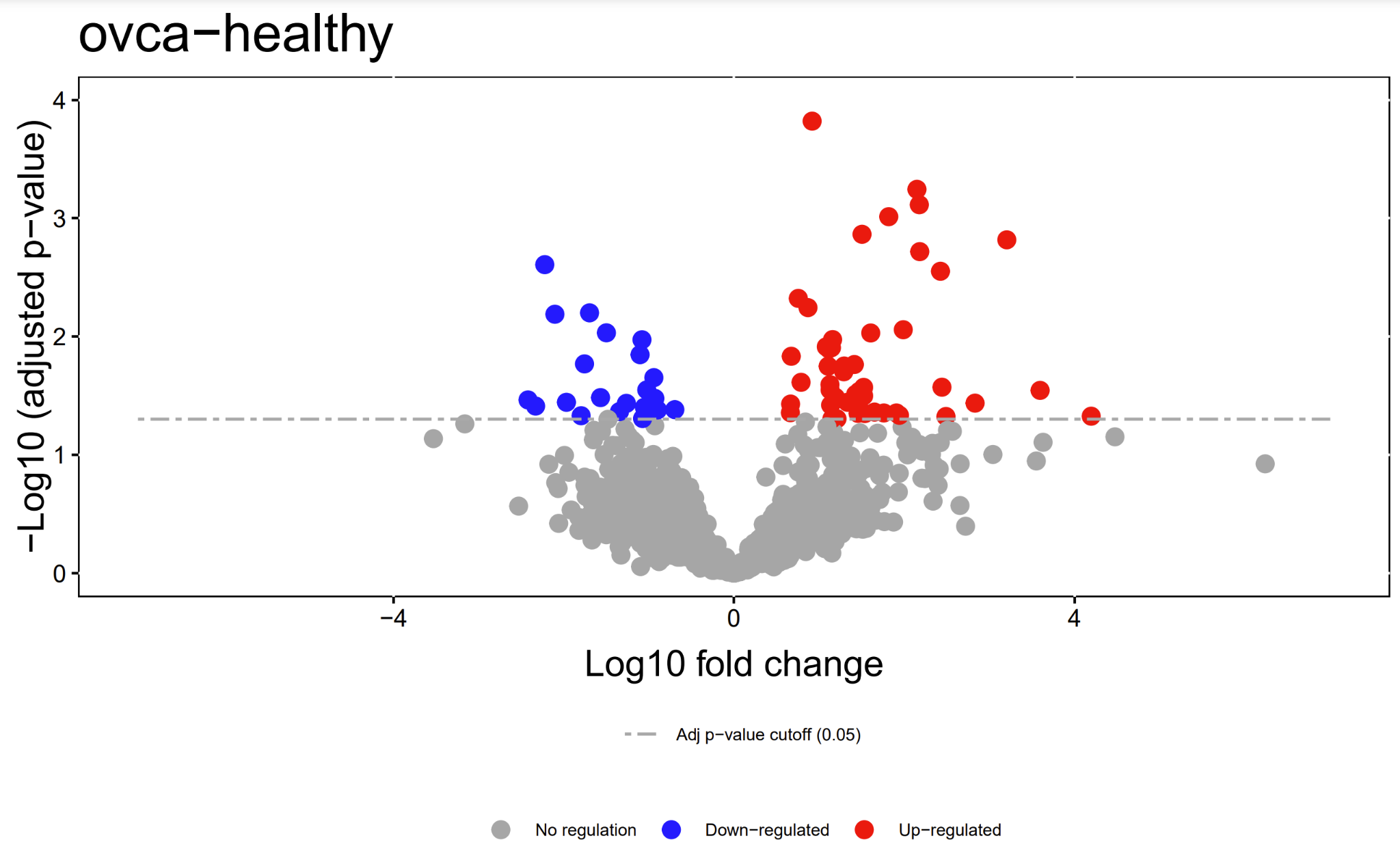

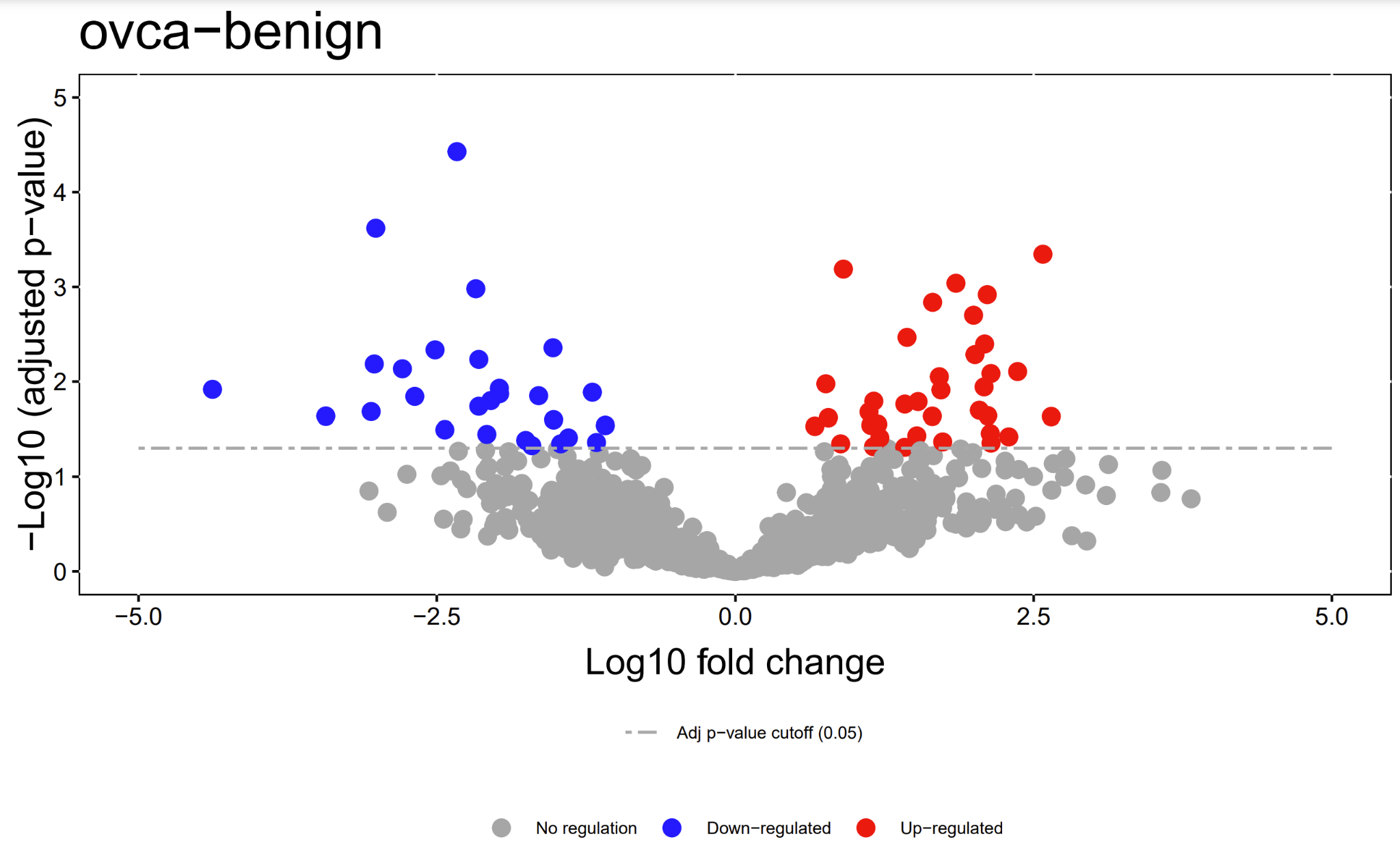

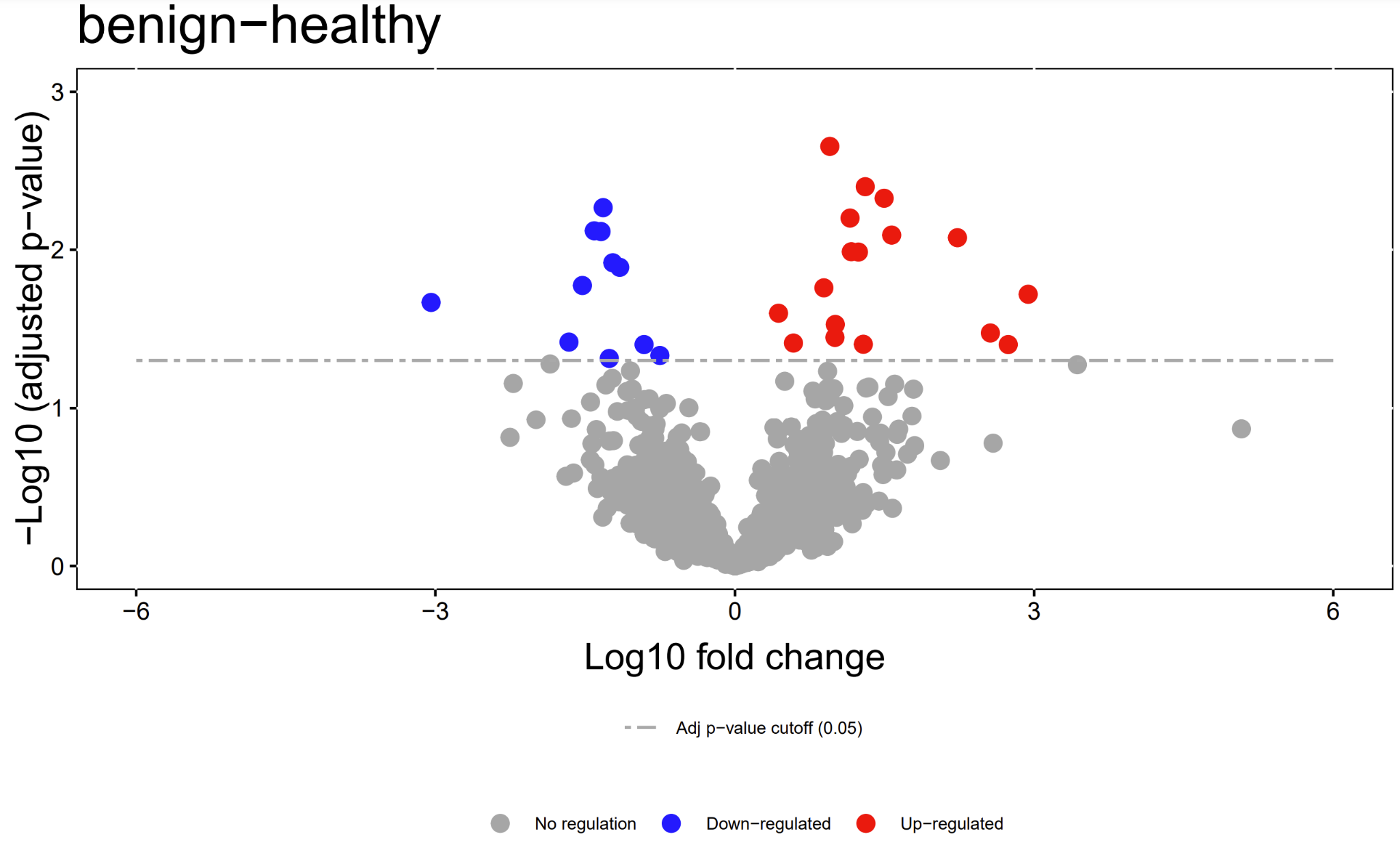

**Figure S3. Human Protein Volcano Plots.** Examples of volcano plots illustrating the log10 fold changes and P-values for comparisons (ovca–healthy, ovca–benign, benign–healthy) for quantified human proteins.
